# Adding insult to injury: Effects of chronic oxybenzone exposure and elevated temperature on two reef-building corals

**DOI:** 10.1101/2019.12.19.882332

**Authors:** Tim Wijgerde, Mike van Ballegooijen, Reindert Nijland, Luna van der Loos, Christiaan Kwadijk, Ronald Osinga, Albertinka Murk, Diana Slijkerman

## Abstract

We studied the effect of chronic oxybenzone exposure and elevated temperature on coral health. Microcolonies of *Stylophora pistillata* and *Acropora tenuis* were cultured in 20 flow-through aquaria, of which 10 were exposed to oxybenzone at a field-relevant concentration of ~0.06 μg L^−1^ at 26 °C. After two weeks, half of the corals experienced a heat wave culminating at 33 °C. All *S. pistillata* colonies survived the heat wave, although heat reduced growth and zooxanthellae density, irrespective of oxybenzone. *A. tenuis* survival was reduced to 0% at 32 °C, and oxybenzone accelerated mortality. Oxybenzone and heat significantly reduced photosynthetic yield in both species, causing a 5% and 22−33% decrease, respectively. In addition, combined oxybenzone and temperature stress altered the abundance of five bacterial families in the microbiome of *S. pistillata*. Our results suggest that oxybenzone adds insult to injury by further weakening corals in the face of global warming.

**Highlights:**

➢ Chronic effect study on corals combining oxybenzone and elevated temperature
➢ Oxybenzone affected photosystem II of coral photosymbionts and altered coral microbiome
➢ Temperature effects were stronger than oxybenzone effects
➢ Sensitivities were species-dependent
➢ Oxybenzone adds insult to injury by weakening corals in the face of global warming

## Introduction

Coral bleaching due to global warming is currently the largest threat to tropical coral reefs (Hughes et al., 2017). Mass bleaching has devastating effects on coral reefs and their ecosystem services, which include coastal protection, fisheries and tourism (Moberg and Folke, 1999). Coral bleaching occurs when seawater temperatures exceed a certain threshold, and involves the expulsion of symbiotic zooxanthellae, which inhabit the coral’s gastroderm (Kleppel et al., 1989; Gates et al., 1992; Iglesias-Prieto et al., 1992; Fitt et al., 2001). Corals do not survive long whilst in a bleached state, most likely due to energy limitation in the absence of photosynthesis (Glynn et al., 1984; Berkelmans et al., 2004).

Next to climate change, seawater pollution has been implicated in coral bleaching. Recently, Brown (2000) discussed the possibility that seawater warming and exposure to pollutant exposure may act additively or synergistically in the process of bleaching. Some explicit examples have been reported since. Morris et al (2019) stated that although coral bleaching is predominantly attributed to photo-oxidative stress, nutrient availability and metabolism also underpin the stability of the coral-zooxanthellae symbiosis. Wooldridge and Done (2009) showed that corals growing in nutrient-enriched coastal waters had a decreased bleaching resistance compared to reefs in oligotrophic oceanic waters. In their study area, eutrophication effectively lowered the bleaching threshold by 1.0–1.5 °C, possibly due to elevated zooxanthellae densities and subsequent higher production of reactive oxygen species. Lapointe et al. (2019) showed that an altered N:P stoichiometry can intensify the effects of high temperature on coral bleaching. Although herbicide levels were not field-relevant, Amid et al. (2018) observed combined effects of elevated temperature and herbicides on the photosynthetic capacity of *Acropora formosa*, indicating the relevance of studying multiple stressors simultaneously. Although these studies provide several examples of additive stress resulting in coral bleaching, research on the combined effects of water quality and water temperature on coral health remains limited (Ban et al., 2014).

An increasing number of studies report the world wide occurrence of synthetic UV filters in coastal waters (Downs et al., 2016; Tsui et al., 2017; Schaap and Slijkerman, 2018; Mitchelmore et al., 2019) as a consequence of either outdoor tourism or waste water discharge due to use of sunscreens or industrial applications (Giokas et al., 2007). In marine waters, UV filters such as oxybenzone are generally found at levels in the order of ng L^−1^ (e.g. Tsui et al., 2014; Sánchez Rodrίguez et al., 2015), but incidental concentrations of over 1 mg L^−1^ have been reported (Downs et al., 2016). Oxybenzone is moderately lipophilic (log_10_ K_ow_= 3.79, log_10_ BAF 2.21-3.01, Tsui et al., 2017) and has been shown to bioaccumulate in clams, urchins and prawns, reaching levels up to 100 ng/g dry weight (Ramos et al., 2015). As oxybenzone and other UV filters recently have been found in wild coral tissue samples at levels up to 38.4 ng/g wet weight in Hong Kong (Tsui et al., 2017) and up to 241 ng/g dry weight in coral tissues of Hawaii (Mitchelmore et al., 2019), it is vital that the environmental consequences of UV filters are studied in more detail.

Organic UV filters such as oxybenzone (benzophenone-3) can be harmful to corals and their larvae (Danovaro et al., 2008; Downs et al., 2016; He et al., 2019). Research suggests that oxybenzone might act as an endocrine disruptor during skeletal development (Downs et al. 2016) and that oxybenzone promotes coral bleaching by inducing a viral lytic cycle in zooxanthellae (Danovaro et al., 2008). Downs et al. (2016) reported an LC50 (24 h) of 139 μg L^−1^ for planulae of corals exposed to oxybenzone, and deformity EC20 levels (24 h) of planulae as low as 6.5 µg L^−1^. He et al. (2019) observed significantly increased settlement failure, bleaching and mortality of *Seriatopora caliendrum* larvae and reported a LOEC of 10 µg L^−1^ for BP-1 and BP-8 exposure, both being degradation compounds of oxybenzone (Kim and Choi, 2014). They reported that coral microcolonies were more sensitive to oxybenzone than larvae (He et al., 2019).

Although the mechanisms underlying oxybenzone toxicity to corals are still unclear, it may involve formation of tissue damaging reactive oxygen species (ROS) (Hanson et al., 2006; He et al., 2019). This mechanism seems similar to the effects elevated seawater temperatures and high light intensity (Dove et al., 2006). During heat and/or light stress, the core photosynthetic machinery within the zooxanthellae known as photosystem II (PSII) becomes less efficient at utilising light energy (Warner et al., 1996; Hill et al., 2009), which is possibly due to ROS causing damage to PSII (Dove et al., 2006). Oxybenzone exposure may increase the ROS pool within coral tissue and zooxanthellae, aggravating the detrimental effects of elevated temperatures on coral reefs. Although Danovaro et al. (2008) describe the promotion of bleaching by oxybenzone through induction of a viral lytic cycle in zooxanthellae, we observed sudden necrosis in corals after exposure to oxybenzone during preliminary studies in our lab (unpublished). This observation, together with the report of Danovaro et al. (2008), might be indicative of a compromised microbiome. A compromised microbiome may alter the physiology of the coral holobiont. For example, changes in the microbiome can reduce the coral’s ability to fix nitrogen, which could be detrimental as corals usually inhabit oligotrophic reef waters (van Oppen and Blackall, 2019).

While a ban on oxybenzone and similar UV filters has been imposed in several places following the precautionary principle (Sirios, 2019), it is yet unknown to what extent field-relevant oxybenzone concentrations affect coral reefs. Even though several toxicological studies involving oxybenzone show impact on corals (Danovaro et al., 2008; Downs et al., 2016; He et al., 2019), the concentrations applied in those studies are several orders of magnitude larger than found on most reefs. In addition, it is unclear whether oxybenzone exposure acts additively or synergistically with elevated seawater temperatures, as both may result in ROS formation, PSII damage and subsequent coral bleaching.

The aim of this study was to determine the combined effect of elevated seawater temperatures and water pollution with oxybenzone on coral health. We tested the hypothesis that oxybenzone-exposed corals are more sensitive to elevated seawater temperatures compared to controls, expressed by an earlier onset of bleaching and necrosis, reduced growth and an altered microbiome composition. Colonies of two common reef building coral species, *Stylophora pistillata* and *Acropora tenuis* were maintained in flow-through aquaria to assess species-dependent sensitivity for elevated temperature and oxybenzone. Half of the corals were exposed to oxybenzone at a nominal level of 1 µg L^−1^, based on levels reported from coastal waters around the island of Bonaire by Schaap and Slijkerman (2018). After two weeks of oxybenzone exposure, half of the controls and half of the oxybenzone-exposed corals experienced a heat wave culminating at 33 °C. During this 6-week study, we monitored survival, growth rate, and photosynthetic efficiency as effective and maximum PSII yield. After 6 weeks, zooxanthellae densities of *S. pistillata* were scored to determine the extent of coral bleaching. In addition, we used 16S rRNA gene sequencing to determine changes in the coral microbiome in response to heat and oxybenzone stress, as the microbiome has been recognized as an essential part of a healthy coral holobiont (van Oppen and Blackall, 2019).

To our knowledge, this study is the first to report on the combined effects of chronic oxybenzone exposure at field-relevant levels and elevated temperatures on coral microcolonies. This study contributes to a better understanding of the interplay between oxybenzone exposure and global warming in terms of coral health, allowing policy makers to re-evaluate the use of UV filters in sunscreens and industrial products.

## Materials and Methods

### Coral husbandry and fragmentation

Captive-bred corals were obtained from Burgers’ Zoo BV (Arnhem, The Netherlands) and cultured at Wageningen University (Wageningen, The Netherlands). The experiment was also conducted at Wageningen University. No approval from an ethics committee was required as scleractinian corals are exempted from legislation concerning the use of laboratory animals in the European Union (Directive 2010/63/EU).

For this study, the Indo-Pacific scleractinian corals *Stylophora pistillata* (Esper, 1797) and *Acropora tenuis* (Dana, 1846) were used. For each species, coral fragments (N◻=◻20 and N=60 for *S. pistillata* and *A. tenuis*, respectively) were cut from a single parent colony and vertically glued onto 5×5 cm PVC tiles (Wageningen UR, The Netherlands) using two-component epoxy resin (Tunze Aquarientechnik GmbH, Germany). Only the growing tips were cut, resulting in uniform fragments of roughly 1 cm in length. All fragments were allowed to recover for 4 weeks in a 550 L holding aquarium before the onset of the 6-week experiment. The holding aquarium was provided with full spectrum white light at an irradiance of 300 μmol m^−2^ s^−1^ (12 h:12 h light:dark regime), created by three 190W LED fixtures (CoralCare, Philips Lighting, The Netherlands). Water flow was provided by several Turbelle nanostream 6085 circulation pumps (Tunze Aquarientechnik GmbH, Germany). Artificial seawater (ASW) for the holding aquarium was prepared using Zoo Mix artificial sea salt (Tropic Marin GmbH, Germany) and deionised water. This mixture was aerated at least 48 hours prior to use. Salinity was kept at 35 g L^−1^ (psu), water temperature at 26 °C, pH at 7.8-8.2, alkalinity at 2.5 mEq L^−1^, calcium at 380-420 mg L^−1^, nitrate at 0-0.25 mg L^−1^ and phosphate at 0-0.05 mg L^−1^. Salinity was measured with a conductivity meter (WTW, Germany), alkalinity was measured with a titralab AT1000 titrator (Hach, Germany). Calcium was measured with an EDTA titration kit (Salifert, The Netherlands). Nitrate and phosphate were measured with colorimetric kits (Red Sea, Israel). The pH was monitored with a pH controller (Resun, China). Every week, 20% of the aquarium water was exchanged to maintain sufficient trace element concentrations. Corals were fed daily with newly hatched brine shrimp (*Artemia salina*), at an end concentration of approximately 250 individuals L^−1^. The aquarium was kept free of algae with five herbivorous fish (two *Zebrasoma flavescens*, one *Acanthurus triostegus*, one *Ctenochaetus strigosus* and one *Siganus vulpinus*), and free of anemones with one *Chaetodon auriga*.

### Experimental setup

Figure 1 shows the experimental setup in which a 2 × 2 experimental design was used, with two temperature regimes (heat wave and control), with and without oxybenzone exposure. To prevent pseudoreplication, the setup contained 5 independent 12L flow-through aquaria for each of the 4 experimental treatments, totalling 20 aquaria. For each temperature regime, one basin with 10 aquaria was supplied with a 300W heater connected to a digital thermostat (Conrad, The Netherlands). To create the heat wave, the digital thermostat of one basin was increased stepwise over a period of three weeks, according to Figure S1, with water temperatures rising from 26 °C to 33 °C. The other basin was kept at 26 °C throughout the experiment. Two 8000 L/h circulation pumps (Tunze, Germany) in each basin prevented the formation of a temperature gradient between the individual aquaria. Within each aquarium, a circulation pump (300 L/h, Eheim, Germany) was placed as well. The light regime was kept the same as in the holding tank, here created by eight 190W LED fixtures (CoralCare, Philips Lighting, The Netherlands). The LED lights were dimmed to 80% of their maximum power and adjusted in such a way that irradiance differences between the 20 aquaria were ± 30 μmol m^−2^ s^−1^. The same ASW as used for the holding tank was used for the aquaria, which was again aerated at least 48 hours prior to use. Three *Acropora tenuis* microcolonies were put into each 12L aquarium, as this species was considered to be highly sensitive. Only one *Stylophora pistillata* microcolony was introduced in each aquarium. Pre-aerated ASW was pumped to the 20 aquaria from two separate 225L header tanks, which contained either control or oxybenzone-spiked ASW (see below for details), using peristaltic pumps (Cole-Parmer, USA). As the corals and algae growing in the 12L aquaria consume significant amounts of nutrients, spiking in the header tanks was required to prevent coral bleaching and necrosis due to lack of nutrients. Therefore, inorganic nutrients were added to both header tanks. Sodium nitrate (NaNO_3_, VWR, The Netherlands) and sodium hydrogen phosphate (Na_2_HPO_4,_ VWR, The Netherlands), were added to reach concentrations of 0.5 mg L^−1^ as NO_3_^−^ and 0.05 mg L^−1^ as PO_4_^3-^. Although these values are significantly higher than measured in oligotrophic reef waters, they were chosen based on prior experience with this setup. The pumping rate was 4L per aquarium per day, equalling a 33% daily ASW turnover. The excess water from each 12L aquarium drained into the water basin via a 20 mm PVC bulkhead, each basin having its own drain (Fig. 1). Each aquarium was also fitted with a perforated lid to prevent cross-contamination between aquaria. Colonies were rotated weekly within each aquarium to average out potential local variations in light and flow intensity. Similar to the husbandry tank, corals in each aquarium were fed daily with brine shrimp (*Artemia salina*) at and end concentration of 250 individuals L^−1^. The coral PVC plates were briefly cleaned weekly with a tooth brush in seawater with the same water chemistry (i.e. from the header tanks) to remove algae. During cleaning, separate brushes and containers were used to prevent sample cross-contamination.

**Figure 1.**
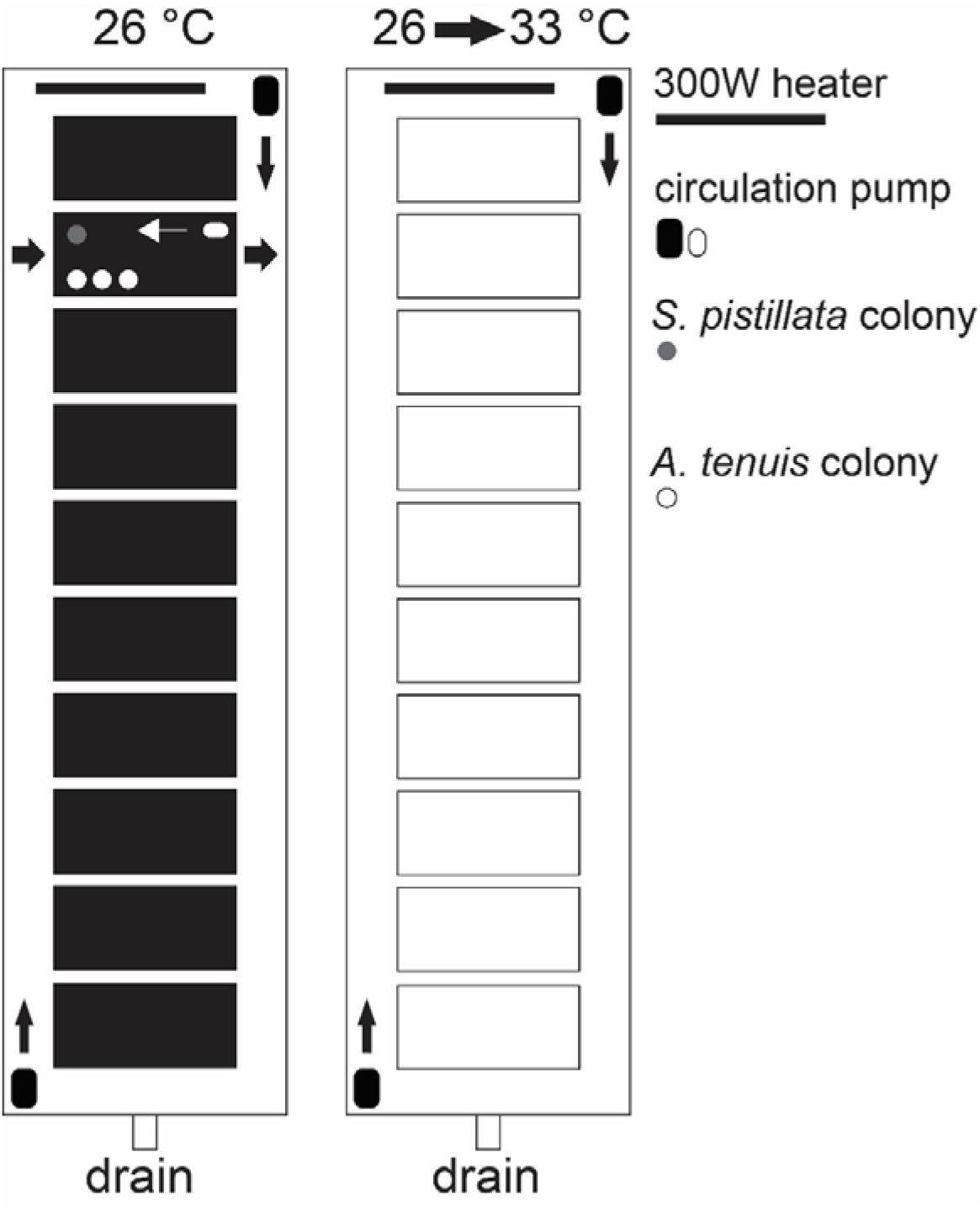
Overview of experimental setup. Thick black arrows indicate seawater flow through an individual 12L aquarium present in a water basin, white arrow shows water flow within the aquarium. Thin black arrows depict circular water flow through the basins, to prevent temperature gradients. White and grey circles depict *S. pistillata* and *A. tenuis* microcolonies, respectively. For clarity, details of only one tank are shown. Control and oxybenzone aquaria were alternated.

### Oxybenzone dosing and recovery

Two 225L header tanks were used to supply the 20 aquaria with ASW, with the control header tank containing ASW + 0.01 mL DMSO L^−1^, and the oxybenzone header tank containing ASW + 0.01 mL DMSO L^−1^ + 1 μg oxybenzone L^−1^. The oxybenzone stock solution was made by dissolving 10 mg oxybenzone salt (Sigma-Aldrich, USA) in 100 mL DMSO (100 mg L^−1^). DMSO was used as the solvent because oxybenzone poorly dissolves in deionised water, at an end concentration (0.001% v/v) far below toxic levels (Galvao et al., 2014). The DMSO-oxybenzone stock solution was dosed to the oxybenzone header tank every other day, when a new batch of ASW was made to maintain sufficient seawater in both header tanks. The DMSO-oxybenzone stock was freshly made from oxybenzone salt every other day and completely shielded from light, to prevent degradation of the oxybenzone stock throughout the experiment.

To determine the actual oxybenzone concentration in the experimental aquaria, water samples were taken twice a week. For each temperature group, 80 mL seawater per aquarium was sampled and combined into one 400 mL sample. Every other week, two control aquaria of each temperature group were sampled as well. Samples were analysed with liquid chromatography (reversed phase Atlantis T3 C18 column) coupled by mass spectrometry (Agilent Zorbax Eclipse XDB-C18 2.1 × 150 mm 3.5µ m column) by Wageningen Marine Research in IJmuiden (The Netherlands). The method was previously validated for a salt water matrix with reproducibility of 3.9%, LOQ 14 ng L^−1^ and LOD 7.2 ngL^−1^. Spiked samples were analysed with each set of samples with recoveries between 88 – 128%. Calibration curves consisted of 6 points between 2.5 and 100 ng ml^−1^ with R^2^>0.999. A more detailed description of the analysis methodology is provided in supplementary information 1 (SI-1).

### Effective and maximum PSII yield

Pulse Amplitude Modulation (PAM) fluorometry was used to non-intrusively determine the effective and maximum quantum yield of photosystem II within the zooxanthellae throughout the experiment. A Diving-PAM-II fluorometer (Heinz Walz GmbH, Germany) was used to measure PSII yield on four consistent sides of each coral microcolony. Effective PSII yield was measured five times per week under light conditions, between 10:30h and 11:30h in the morning, whereas maximum PSII yield was measured once per week in darkness between 7:30h and 8:30h. Measuring and saturating lights were provided by a full spectrum lamp within the fluorometer and delivered to the microcolonies via a 5 mm diameter fibre optic cable. The fibre optic was positioned approximately 2 mm from the surface of the coral fragment. Under dark conditions, minimum (F_0_) and maximum fluorescence (F_m_) were measured. Under light conditions, F and F_m’_ were measured. These parameters were used to calculate effective (ΔF/F_m_’) and maximum (F_v_/F_m_) PSII yield (Schreiber et al., 1986).

### Specific growth rate

To determine coral growth rates, the buoyant weighing method (Davies, 1989) was used. Buoyant weight was measured at the start of the experiment by suspending each coral from a hook tied to a nylon string in a defined seawater volume at constant depth. The hook was attached to an under-weighing analytical balance with 0.1 mg accuracy (Ohaus, Germany) using the nylon string (Osinga et al., 1999). All coral microcolonies were weighed before and after mounting on a plate with two-component epoxy resin, allowing for calculation of net coral weights throughout the experiment. A round metal cup was used to suspend the coral from the hook before it was glued to the PVC plate. The plates had a perforation to allow them to be suspended on the hook after the corals were glued. Seawater was maintained at 26°C and 35 g L^−1^ (psu) salinity to maintain constant density. Based on the branching nature of the coral species used, exponential growth was assumed and specific growth rates (μ) were calculated using the formula:

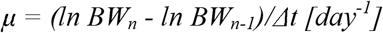

where μ is the specific growth rate (day^−1^), BW_n_ is buoyant weight at the end of a growth interval, BW_n-1_ is buoyant weight at the start of a growth interval and Δt is the time in days between measurements. To determine growth rates before and during the heat wave, all corals were weighed on day 0, 16 and 41, where the latter two days correspond to temperatures of 26°C and 33 °C for the heat wave group, respectively.

### Zooxanthellae density

At termination of the experiment (41 days), fragments (N=5 per treatment, N=20 in total) of approximately 5 mm were cut from each *S. pistillata* microcolony with disinfected pliers and placed on Kimwipes (Sigma-Aldrich, USA) in a fume hood. The length and width of each fragment were measured with a ruler to allow calculation of tissue surface area, where a fragment was considered to be a semi-open cylinder, resulting in the following formula: 2*π*r*h + π * r^2^. Here, *r* is the sample radius, and *h* is the sample height in cm. Subsequently, coral fragments were put inside 15 mL tubes using sterile tweezers and coral tissue was removed from the calcium carbonate skeleton using an air gun during 3 minutes. Tubes were sealed with parafilm to avoid splashing of tissue from the tube. After tissue removal, the bare coral skeleton was removed from the tube, 10 mL ASW (35 g L^−1^) was added and each sample was briefly shaken for 10 seconds. When all samples were processed, all tubes were shaken for another 3 minutes to dissolve all tissue. Next, samples were centrifuged at 2000 *g* for 10 minutes, in order to separate coral tissue from zooxanthellae. After centrifugation, the supernatant of each sample was discarded and zooxanthellae pellets were homogenised in 750 μL of ASW. After determining the volume of each sample using a 1000 μL pipet, 2 subsamples were transferred to a Fuchs-Rosenthal counting chamber to measure zooxanthellae concentrations. Finally, the volume and zooxanthellae concentration of each sample were multiplied and divided by the surface area of each fragment. Zooxanthellae densities were expressed as cells per cm^2^ coral tissue.

### DNA isolation, PCR and sequencing

After zooxanthellae counting, the remainders of the tissue samples were further processed for microbiome analysis by 16S rRNA gene sequencing. All samples were centrifuged at 5000 g for 15 minutes to pellet zooxanthellae and bacteria. Supernatants were discarded, and DNA was extracted and purified from the pellets using the DNeasy Blood & Tissue kit (Qiagen, Hilden, Germany), with pre-treatment for gram-positive bacteria according to the manufacturer’s protocol. For each sample and corresponding negative controls, 16S rRNA genes were amplified by PCR using the following primers: 27F_BCtail-FW (TTTCTGTTGGTGCTGATATTGC_AGAGTTTGATCMTGGCTCAG) and 1492R_BCtail-RV (ACTTGCCTGTCGCTCTATCTTC_CGGTTACCTTGTTACGACTT), each containing an 5’ extension allowing for subsequent barcoding by PCR, and Phire Tissue direct PCR master mix (Thermo Fisher) using the following amplification cycles: 3 min at 98 °C, 30 cycles of 8s at 98°C; 8s at 58°C; 30s at 72 °C, final extension of 3 min at 72 °C. Obtained amplicons were purified using the QIAquick PCR purification kit (Qiagen, Hilden, Germany). Amplicons for each sample were barcoded using the Oxford Nanopore “PCR Barcoding Expansion Pack 1-96 (EXP-PBC096)”, and sequenced on the MinION according to manufacturers “1D PCR barcoding (96) genomic DNA (SQK-LSK108)” protocol (Oxford Nanopore Technologies).

### Microbiome analysis

A total of 572,828 reads were obtained, of which 201,065 reads were successfully classified and passed quality filtering (Q score threshold ≥8). The Fastq 16S workflow revision 3.2.0 with the 16S classification version 3.0.0 were used on the Oxford Nanopore Epi2me platform (Metrichor, Oxford, UK; https://epi2me.nanoporetech.com) to classify raw sequences. A taxonomic hierarchy of all the classified taxa was obtained using the Entrez Taxonomy NCBI database and the taxize package in R (Chamberlain and Szocs, 2018), which was used to generate an OTU table on order, family and genus level. Data visualisation and statistical tests were performed using the vegan (Oksanen et al., 2018), ggplot2 (Wickham, 2016) and phyloseq (McMurdie and Holmes, 2013) packages. Alpha diversity (within-sample diversity) was calculated using two indices: observed OTU richness and ACE (abundance-based coverage estimator). To test for differences in alpha diversity between temperature and oxybenzone treatments, a generalised linear model with a negative binomial distribution was fitted to the data. To assess differences in beta diversity (between-sample diversity) with temperature and oxybenzone treatments, Bray-Curtis (Bray and Curtis, 1957) dissimilarities were calculated. These metrics were then visualised with a Principal Coordinates Analysis (PCoA) plot and compared in a permutational analysis of variance (PERMANOVA) with 9,999 permutations. All data conformed to the underlying assumptions of PERMANOVA, with multivariate spread being equal among groups (similar to homogeneity of variances in an ANOVA, Anderson 2017). Families that showed significant differences in abundance between temperature and oxybenzone treatments were determined using the DESeq2 package (Love et al., 2014) with Benjamini-Hochberg corrected *p*-values.

### Statistical analysis of PSII yield, coral growth and zooxanthellae density

One aquarium was considered the experimental unit, therefore data obtained from multiple *Acropora tenuis* microcolonies within one aquarium were pooled. As only one *Stylophora pistillata* microcolony was put into each aquarium, no pooling was required for this species. Normality of data was first tested by plotting residuals of each dataset versus predicted values, and by performing a Shapiro-Wilk test. Homogeneity of variances was determined using Levene’s test. All data were found to be normally distributed and showed homogeneity of variances (*p*>0.050) after arcsine transformation (for PSII yield). When the assumption of sphericity, tested with Mauchly’s test, was violated, degrees of freedom were adjusted using Greenhouse-Geisser. We used a three-way mixed factorial ANOVA to test the effects of temperature, oxybenzone exposure and time on effective and maximum PSII yield of *S. pistillata* and *A. tenuis*, with temperature and oxybenzone as between-subjects factors and time as within-subjects factor. Simple effects analysis was used to break down interactive effects. A two-way factorial ANOVA was used to determine the effects of temperature and oxybenzone exposure on specific growth rate and zooxanthellae density of *S. pistillata*. Due to lack of surviving *A. tenuis* samples in the heat wave group at the end of the experiment, data for SGR and zooxanthellae density could not be analysed for this species. A *p*<0.050 value was considered statistically significant. Statistical analysis was performed with SPSS Statistics 25 (IBM, Somers, USA). Graphs were plotted with SigmaPlot 12 (Systat software, San Jose, USA). Data presented are expressed as means ± standard error of the mean (s.e.m.), unless stated otherwise.

## Results

### Water chemistry

Water chemistry parameters (salinity, alkalinity and calcium levels) remained stable throughout the study period and were highly similar among treatments (Fig. S1). Ammonium, nitrate and phosphate were consistently below detection levels (<0.01 mg L^−1^, not shown). Measured oxybenzone concentrations were 0.05±0.03 μg L^−1^ for the 26 °C aquaria and 0.06±0.05 μg L^−1^ for those experiencing a heat wave, thus comparable between treatments. Some variation over time was observed (Fig. S1).

### Main effects of elevated temperature and oxybenzone

The main effects of oxybenzone exposure and the elevated temperature on coral health and growth are summarized in Table 1.

**Table 1.**
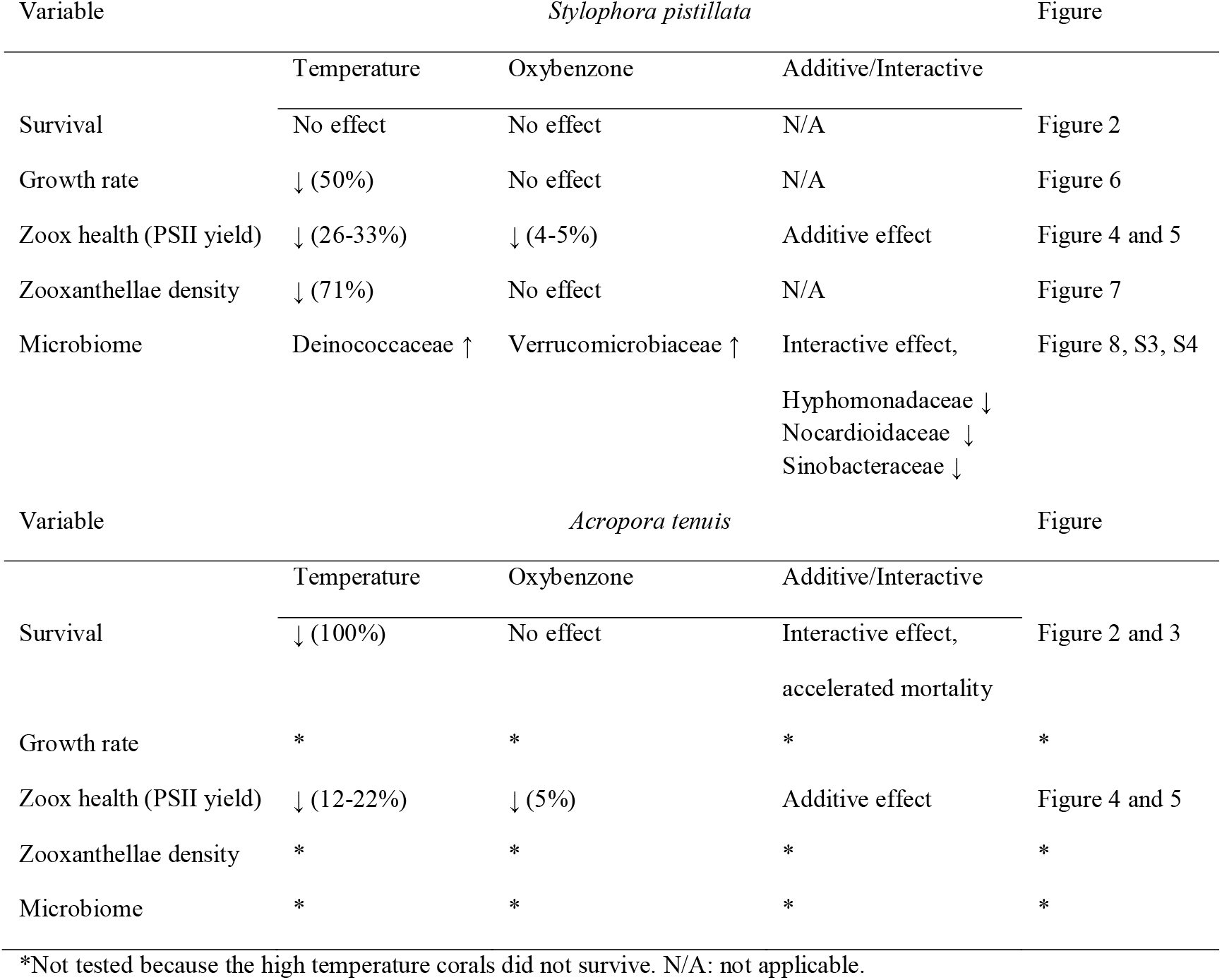
Overview of observed effects of chronic oxybenzone exposure and the heat wave on two reef-building coral species.

All *S. pistillata* microcolonies survived the experiment. At the end of the experiment (Day 41), *S. pistillata* colonies growing at 33 °C exhibited signs of bleaching (Fig. 2A). A distinct green colouration at the tips became apparent, probably due to green fluorescent protein production. Survival of *A. tenuis* stabilised at 67% for the 26 °C treatment without oxybenzone (Fig. 3). High temperatures markedly affected *A. tenuis* survival, with corals exposed to the heat wave without oxybenzone exhibiting a gradual decrease in survival rate over time and a steep decline towards 0% survival at 32 °C, between Day 32 and 33 (Figs. 2B and 3).

**Figure 2.**
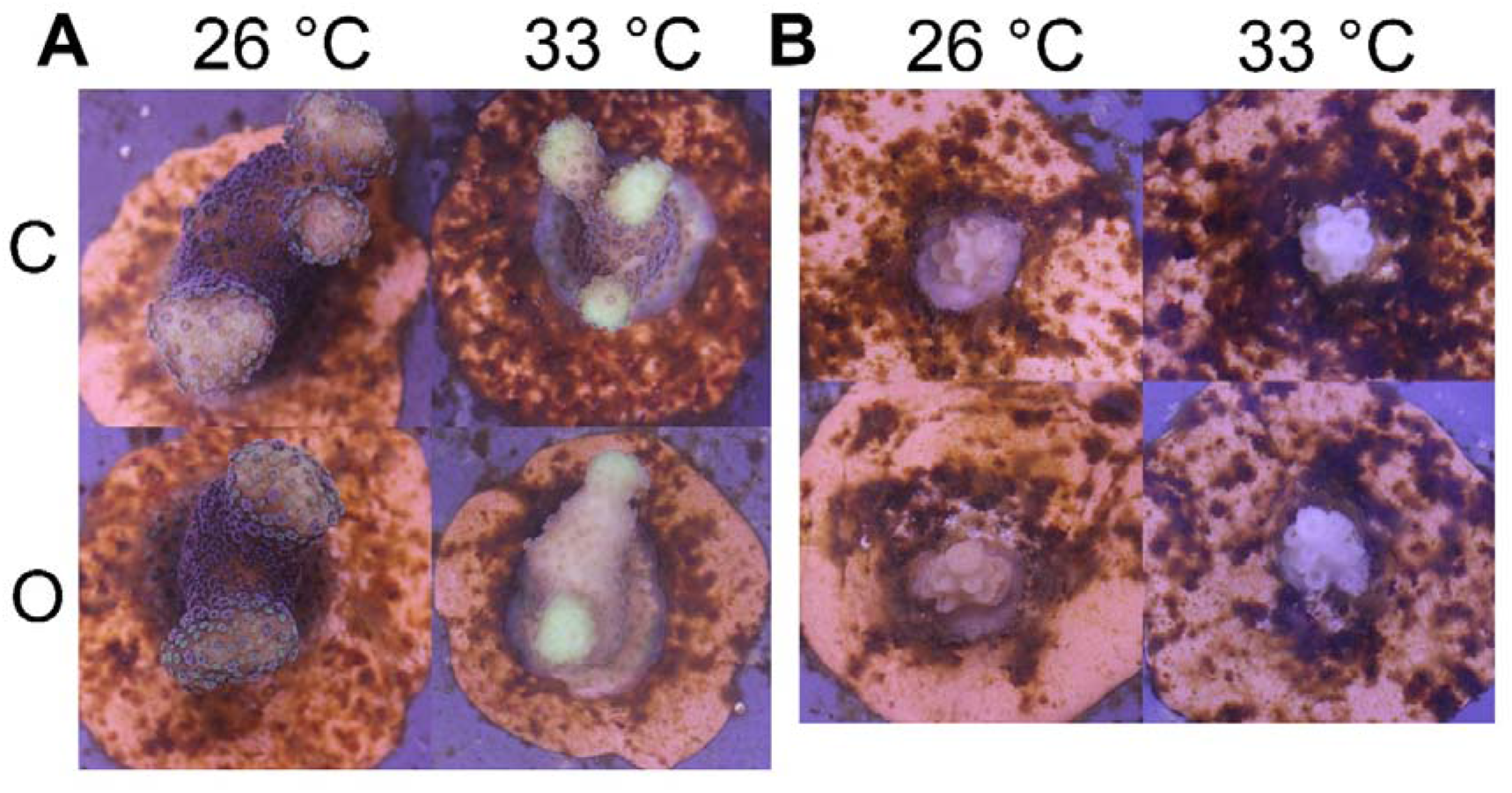
Representative photographs of *S. pistillata* (left group, clear bleaching at 33 °C) and *A. tenuis* (right group, 100% mortality already at 32 °C) microcolonies at the end of the experiment (day 41). C: control, O: oxybenzone. Black scale bars: 5 mm.

**Figure 3.**
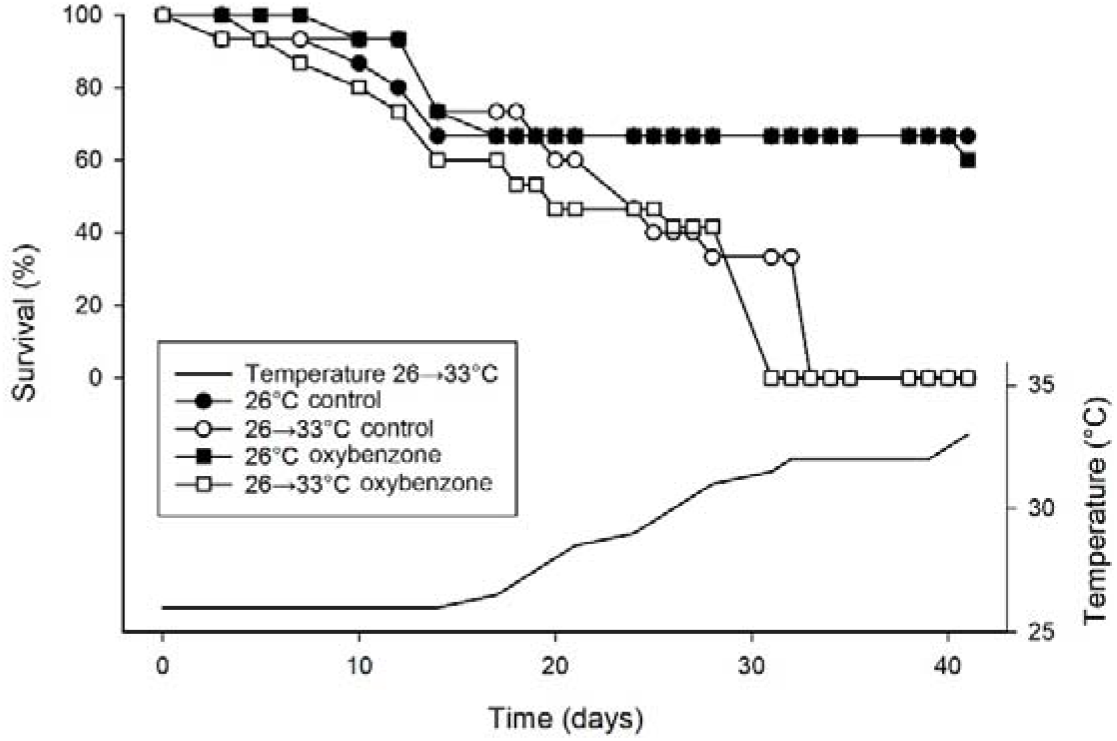
Decreasing survival of *A. tenuis* microcolonies during the experiment with constant (solid symbols) or increasing temperature (open symbols). Circles indicate control aquaria, −1 squares depict aquaria with oxybenzone (~0.06 μg L^-1^). All *S. pistillata* microcolonies survived the experiment and are therefore not shown.

In the 26 °C treatments, the effective and maximum PSII yield were constant for both coral species over the full experimental period of 41 days (Figs. 4 and 5). Temperature increase negatively affected effective PSII yield in both species. For *S. pistillata*, an interactive effect of temperature treatment and time on effective PSII yield was found (Fig. 4A, Table S1). Up to Day 24 (29 °C), PSII yield remained constant (simple effects, *p*>0.05). However, from Day 31 onwards (31.5 °C), a consistent reduction in effective PSII yield was detectable, with a 33% decrease at Day 41 as compared to the control temperature (simple effects, *p*<0.05, Fig. 4A). The 26 °C treatment showed a consistent yield over time, with no temperature effect (simple effects, *p*>0.05, Fig. 4A). For *A. tenuis*, no statistical effect of temperature could be calculated as the corals died during the heat wave. The data, however, show that effective PSII yield of the last surviving corals was reduced by 22% at Day 32 (32 °C, Fig. 4B).

**Figure 4.**
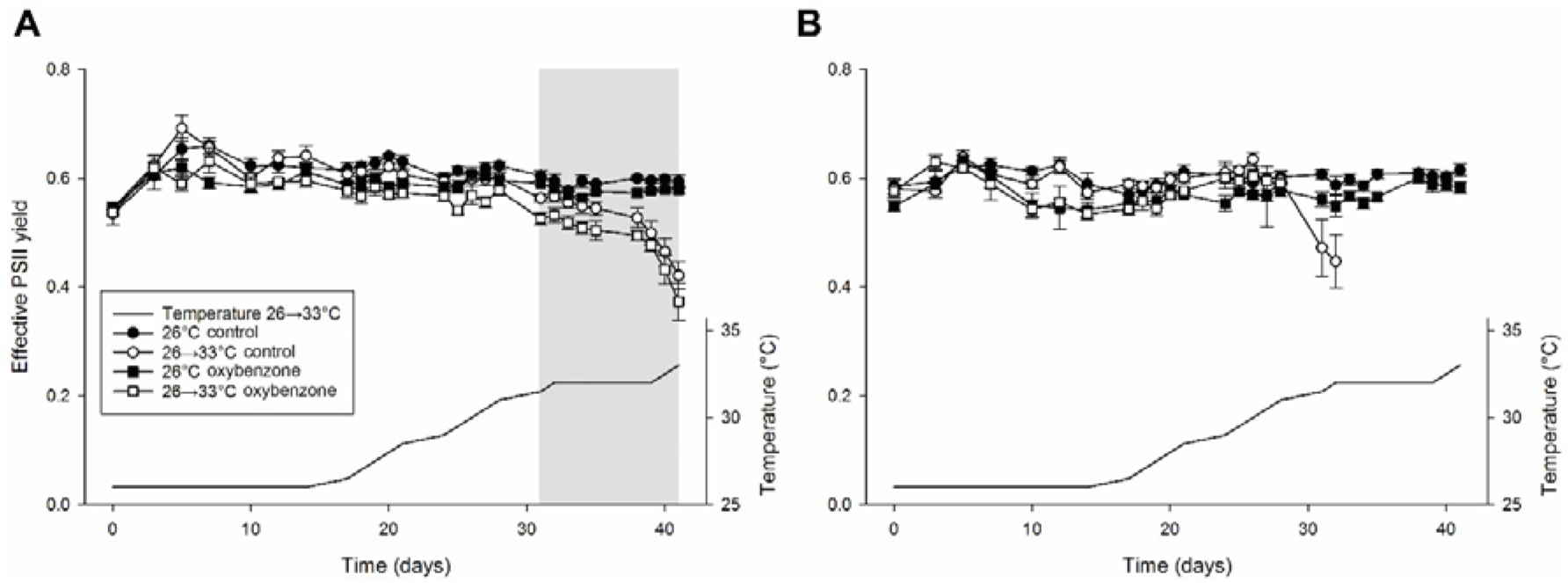
A: Effective photosystem II (PSII) yield of *S. pistillata* microcolonies over time with constant or increasing temperature. Grey area indicates significant effect of temperature (*p*<0.050) in the period spanning Day 31 to 41 of the experiment (N=5). B: Effective PSII yield of *A. tenuis* microcolonies over time (N=2-5). The effect of temperature is not displayed as it could not be calculated due to coral mortality. Values are means ± s.e.m.

**Figure 5.**
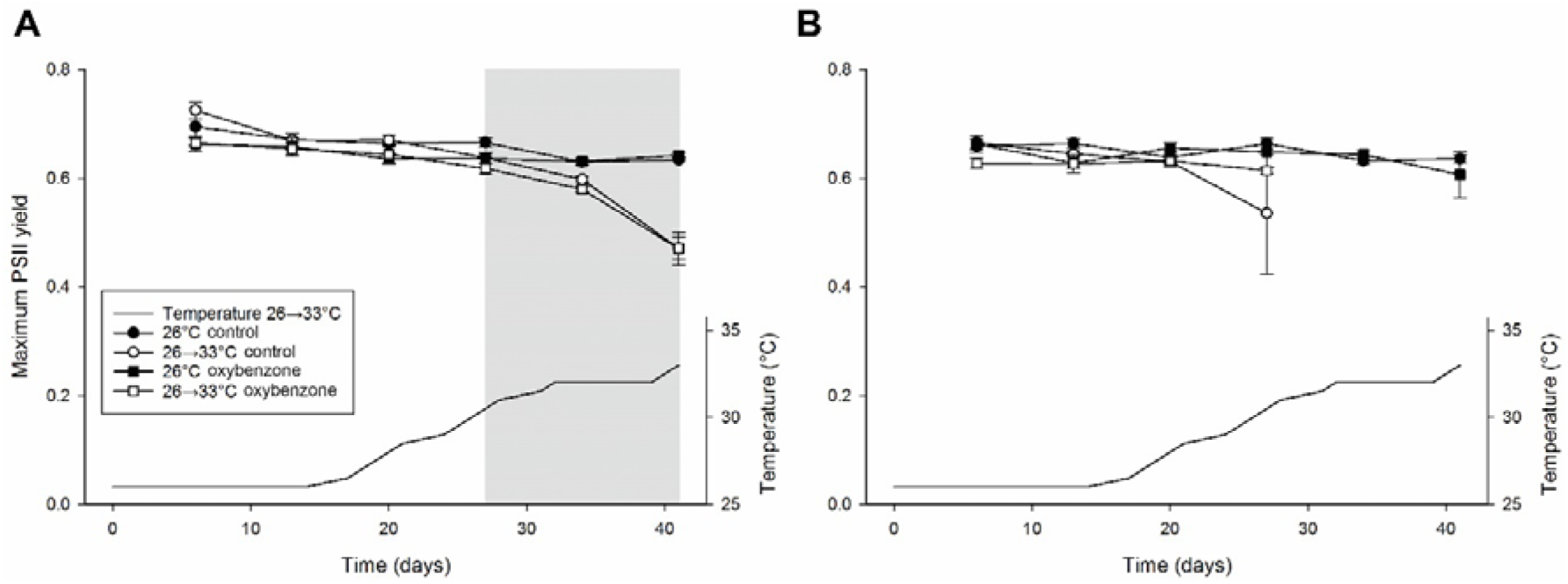
A: Maximum photosystem II (PSII) yield of *S. pistillata* microcolonies over time. Grey area indicates significant effect of temperature (*p*<0.05) in the period spanning Day 27 to 41 of the experiment, irrespective of oxybenzone (N=5). B: Maximum PSII yield of *A. tenuis* microcolonies over time (N=2-5). The effect of temperature is not displayed as it could not be calculated due to coral mortality. Values are means ± s.e.m.

The effect of temperature treatment on maximum PSII yield of *S. pistillata* was also found to increase in time (Table S1). Up to Day 20, (28 °C), no decrease in maximum PSII yield was found. However, from Day 27 onwards (30.5 °C), a consistent reduction in maximum PSII yield was detectable, with a decrease of 26% at day 41 as compared to the 26 °C treatment (simple effects, *p*<0.05, Fig. 5A). Again, the 26 °C treatment showed a consistent maximum yield over time, with no temperature effect (simple effects, *p*>0.05; Fig. 5A). For *A. tenuis*, the maximum PSII yield was reduced by 12% at Day 27 (30.5 °C (Fig. 5B), although not significant (Table S1).

The growth of *S. pistillata* microcolonies was inhibited by 50% during the heat wave (Days 16-41, Fig. 6). Zooxanthellae density in *S. pistillata* was reduced by 71% at Day 41 (33 °C, Fig. 7). Zooxanthellae density and growth rate in *A. tenuis* could not be determined because all microcolonies had died during the heat wave.

**Figure 6.**
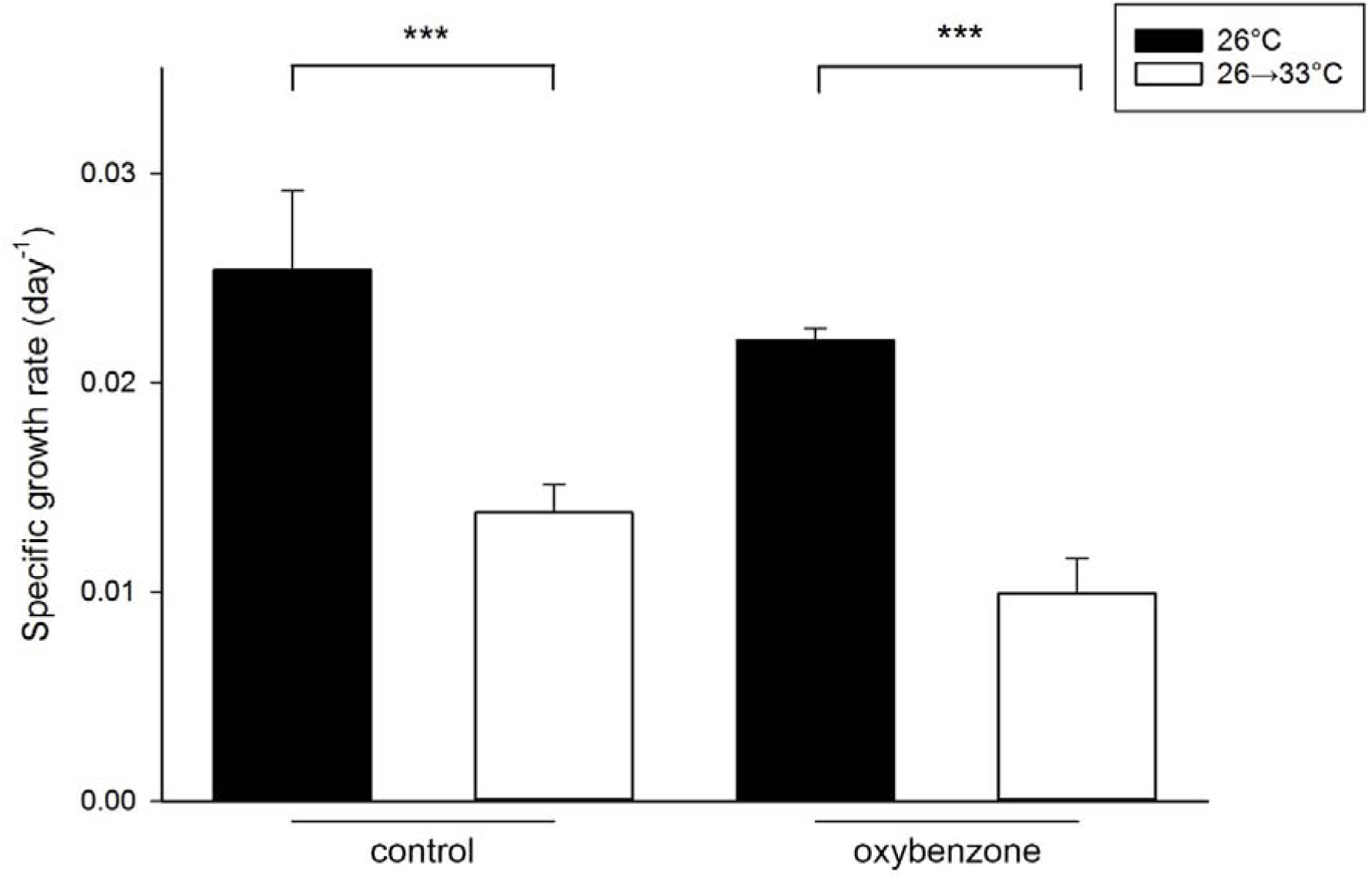
Specific growth rate of *S. pistillata* microcolonies during the heatwave period (day 16 to 41). Asterisks indicate significant effect of temperature (*p*<0.001), irrespective of oxybenzone. Values are means + s.e.m. (N=5).

**Figure 7.**
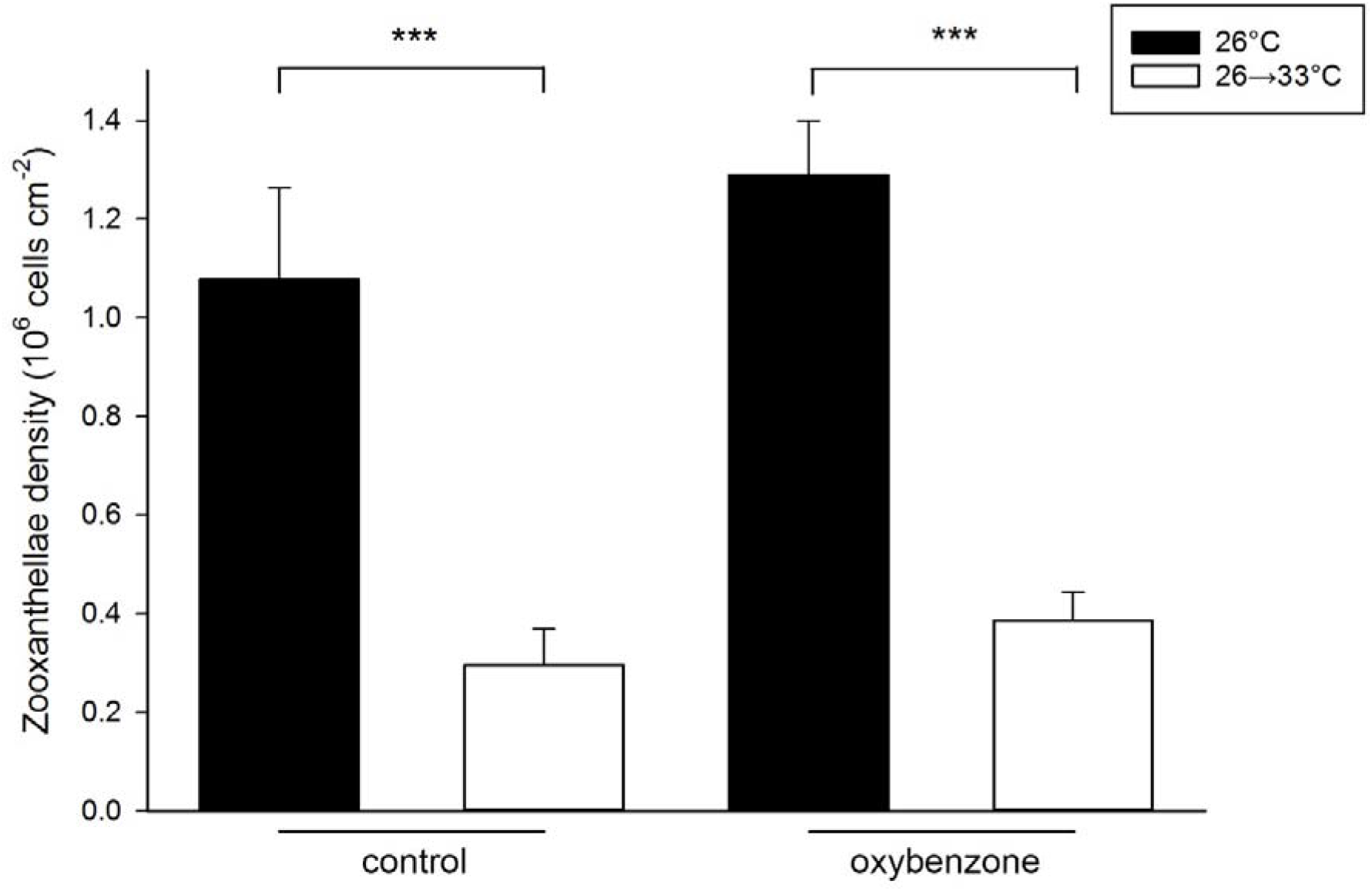
Zooxanthellae density of *S. pistillata* microcolonies at day 41. Asterisks indicate significant effect of temperature (*p*<0.001), irrespective of oxybenzone. Values are means + s.e.m. (N=5).

Only minor effects of oxybenzone exposure alone on both species were observed, for a limited number of parameters. A main effect of oxybenzone on effective PSII yield of both species was found, with a consistent 5% reduction throughout the experiment as compared to controls (Fig. 4, Table S1). Maximum PSII yield was affected by oxybenzone for *Stylophora pistillata* only, with a 4% reduction throughout the experiment (Fig. 5, Table 1).

Oxybenzone did not affect the survival, growth rate and zooxanthellae density of *S. pistillata* (Table S1, Fig. 6–7). Survival of *A. tenuis* at 26 °C was hardly affected by oxybenzone (Fig. 3). For *A. tenuis*, zooxanthellae density and growth rate could not be determined due to observed mortality.

### Combined effects of elevated temperature and oxybenzone

Combined exposure to oxybenzone and elevated temperature accelerated mortality in *A. tenuis* with 2 days compared to temperature elevation alone (Fig. 3). Mortality towards 100% was steep from Day 28 (31 °C) onwards for the oxybenzone-exposed group, compared to Day 32 (32 °C) onwards for the control group. None of the oxybenzone-exposed *A. tenuis* colonies survived at 31.5 °C, whereas the control colonies all died at 32 °C. No interactive effects of oxybenzone and temperature elevation were observed for PSII yield, coral growth or zooxanthellae density (Table S1).

Because of the *A. tenuis* mortality only the microbiome of *S. pistillata* could be analysed. This microbiome contained ~95 genera (mean of observed richness, ranging from 62-155 genera), ~60 families (ranging from 45-86), and ~38 orders (ranging from 31-50). All raw data and a list with the observed taxa have been uploaded and are publicly available in the European Nucleotide Archive. Combined exposure to oxybenzone and elevated temperature resulted in a distinct change in the microbiome (Figs. S3 and S4). Although a principal coordinates analysis (PCoA) based on Bray-Curtis distance at order, family or genus level showed no clear pattern (Fig. S3, family level shown only), a PERMANOVA on microbiome beta diversity revealed a significant interactive effect of temperature and oxybenzone (Table S2). On family level, the relative abundance of the families Hyphomonadaceae, Nocardioidaceae and Sinobacteraceae had declined in the combined stressor exposure (Fig. 8, Table S3). As a result of oxybenzone exposure, the Verrucomicrobiaceae had increased, but only at 26 °C (Fig. 8). Elevated temperature only resulted in increased abundance of Deinococcaceae (Fig. 8).

**Figure 8.**
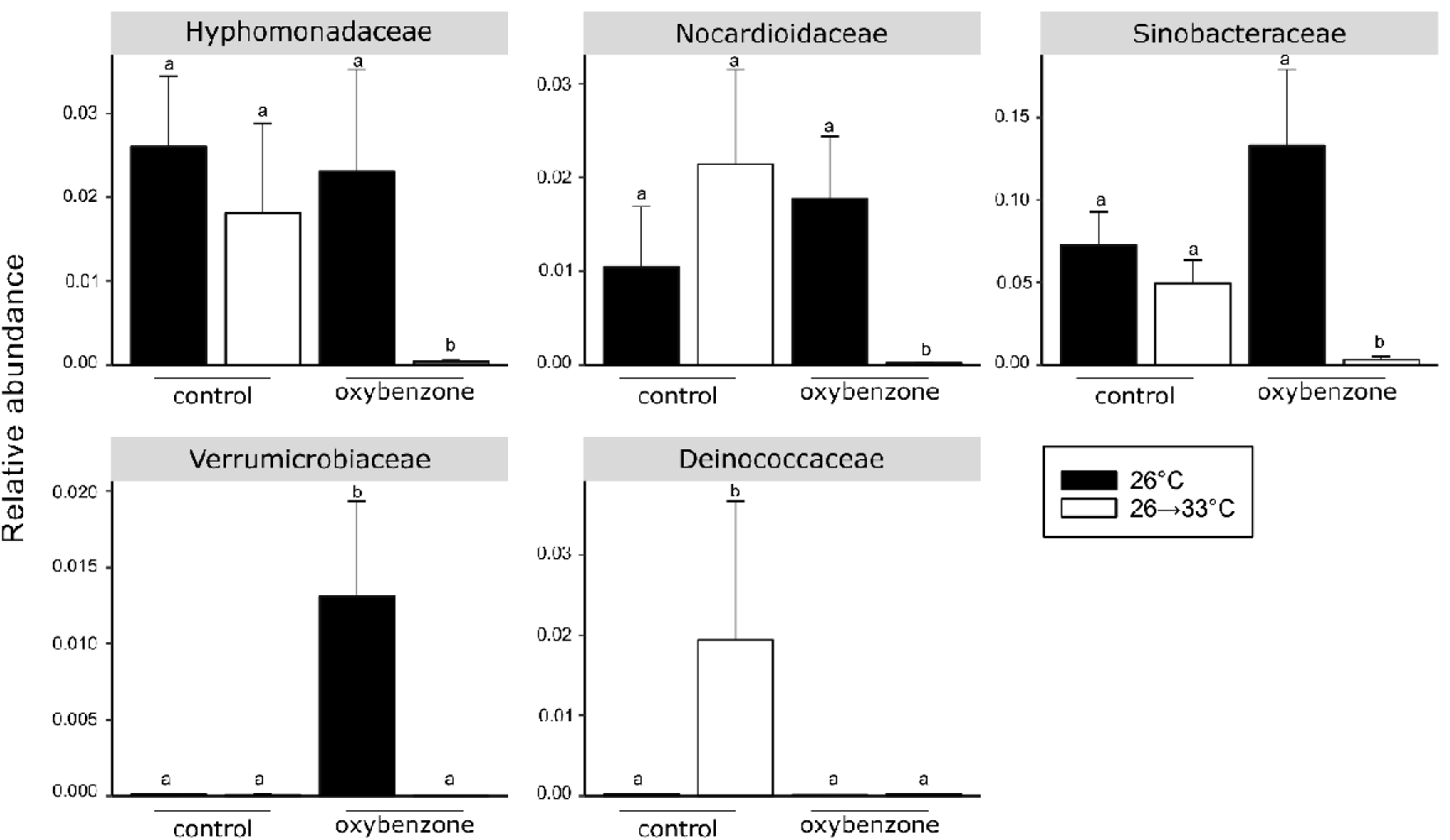
Post-hoc analysis with Benjamini-Hochberg correction of the five families within the *S. pistillata* microbiome which significantly responded to the temperature and oxybenzone treatments. Groups with different superscripts differ significantly (p<0.05). Values are means + s.e.m. (N=5).

On genus level, combined exposure to heat stress and oxybenzone resulted in a similar pattern as seen at family level, with reduced abundance of 10 genera and increased abundance of only one genus (*Marivita*, Figure S4).

## Discussion

The aim of this study was to determine whether elevated seawater temperatures and water pollution with the UV filter oxybenzone act synergistically on endpoints relevant to coral health, in particular to the species *S. pistillata* and *A. tenuis*. The hypothesis is that corals exposed to field-relevant oxybenzone concentrations are more sensitive to elevated seawater temperatures compared to controls, because of a possibly related oxidative stress mechanism. This hypothesis was tested in an experimental setup and is partially supported by the data, which will be discussed in detail below.

### Experimental setup

Seawater chemistry was constant among treatments, with the majority of coral microcolonies in the control group remaining healthy during the full experimental period. The chosen nominal level of oxybenzone was a field-relevant concentration of 1 µg L^−1^ and the re-dosing of oxybenzone was designed to keep the compound concentration within a reasonable range. To check the actual concentrations water samples were analysed weekly. This revealed that the exposure concentration throughout the experiment was much lower than anticipated, being on average 0.06 µg L^−1^. The difference can be explained by the relatively lipophilic nature of oxybenzone with a log_10_ K_ow_ of 3.79 (Tsui et al., 2017). Thus, oxybenzone may have adhered to the plastic tubing used to pump seawater to the aquaria, as well as to the plastic aquaria themselves. Oxybenzone is also taken up by corals (Tsui et al., 2017; Mitchelmore et al., 2019). After uptake by the coral, it is likely that corals metabolize oxybenzone to benzophenone-1 and benzophenone-8 (Kim and Choi, 2014; Tsui et al. 2017). Interestingly, He et al. (2019) found coral species to be more sensitive to BP-8 than to oxybenzone. As we neither measured BP-1 nor BP-8 levels, neither in the aquaria nor in the corals, we cannot exclude that the observed effects in our study resulted from these metabolites. We do show, however, that chronic exposure to an even 20 times lower oxybenzone level (60 ng L^−1^) than originally planned, does induce adverse effects. Field-relevant levels found at locations near coral reefs across the world are in the range of 10 ng L^−1^ with maxima up to 1.5 ug/L (e.g. Tashiro & Kameda, 2013; Tsui et al., 2014; Schaap & Slijkerman, 2018). The adverse effects observed in this laboratory study might thus not be excluded in field situations for which chronic exposure to oxybenzone exists.

### Effects of elevated temperature

Elevated temperature had the most pronounced effects on both coral species and on more end-points (PSII yield, coral growth and zooxanthellae density). The heat wave affected *A. tenuis* more than *S. pistillata*; *A. tenuis* showed 100% mortality, whereas *S. pistillata* only bleached (measured as loss of zooxanthellae), but did not die. The pronounced green colouration of the *S. pistillata* microcolony tips during the heat wave suggests green fluorescent protein (GFP) production, which is known to have antioxidative properties (Bou-Abdallah et al. 2006; Salih et al. 2000) and could help the corals combat the effects of heat and oxybenzone stress. The observed species-dependent thermal sensitivity is in line with previous research (Marshall and Baird, 2000; Loya et al., 2001; Wilkinson, 2004; Guest et al., 2012). Stimson et al. (2002) ranked *Acropora* to be more susceptible to temperature-induced mortality than *Stylophora*, an assessment in line with the observations in our study.

### Effects of oxybenzone

The effects of oxybenzone were statistically significant, but limited and subtle compared to the effects of elevated temperature. Oxybenzone alone hardly affected coral survival, in contrast with He et al. (2019), who reported a bleaching rate of 95% and a mortality rate of 5% in *S. caliendrum* microcolonies at 1000 µg L^−1^ after 7 days of exposure. This discrepancy is likely caused by the much higher concentration (four orders of magnitude) used in their study.

A 4-5% PSII yield reduction due to chronic oxybenzone exposure was found for *S. pistillata* and *A. tenuis*, which could be explained by oxidative damage to zooxanthellar thylakoid membranes (Downs et al. 2016). For *S. pistillata*, oxybenzone also had a more pronounced effect on effective PSII yield (measured in the light) as compared to maximum PSII yield (measured in darkness), possibly due to phototoxicity of oxybenzone (Downs et al. 2016). We did not observe an oxybenzone effect on zooxanthellae density, although a slight positive trend was visible. This is in contrast with Danovaro et al. (2008), who showed viral lysis of zooxanthellae after exposure to 33 uL/L oxybenzone exposure. Similar to the study of He et al. (2019), this observational difference can likely be explained by the much lower applied oxybenzone levels in this study, although differential susceptibility to pollution is known to exist between coral species (Fabricius, 2005; Negri et al., 2011; Downs et al., 2016; He et al., 2019).

The ecological significance of photo-inhibition due to pollution should not be underestimated. Cantin et al. (2007) observed reduced effective quantum yield (20%) in two Acroporids and *Pocillopora damicornis* after chronic exposure to the photosynthesis-inhibiting herbicide diuron at 1 μg L^−1^. These authors also provided evidence for a link between reduced energy acquisition (lower total lipid content) due to PSII photo-inhibition and a 6-fold reduced reproductive output (in terms of polyp fecundity) in zooxanthellate corals in an experimental setting. In our study, oxybenzone exposure at a level of 0.06 µg L^−1^ resulted in a PSII yield reduction of 4-5% in both species. Although we did not assess coral energy acquisition and reproductive output, we cannot exclude a negative long-term impact on metabolism, with decreased growth and reproductive capacity. Although we did not observe a significant effect on coral growth, a trend with 16% growth reduction in oxybenzone-exposed corals was found, possibly caused by PSII inhibition. Significant growth inhibition cannot be excluded at higher oxybenzone levels, longer exposure intervals or more sensitive coral species. The ecological and economic consequences of coral growth inhibition could be severe, as currently some reefs already cannot outgrow sea-level rise, posing a critical threat to the adjacent coastal region (De Bakker et al., 2019). Additional studies should focus on impaired PSII and related growth inhibition by UV filter exposure in more detail in order to evaluate its ecological significance.

### Effects of combined stress

Elevated temperature and oxybenzone exposure each inhibited photosynthetic yield and these effects are additive in both coral species. This is not in line with our hypothesis that oxybenzone aggravates the effects of elevated temperature on coral health. The heat wave, however, did accelerate mortality of *A. tenuis* when oxybenzone was dosed. This is in agreement with our hypothesis and with the findings of Danovaro et al. (2008) who describe stronger oxybenzone effects at higher temperatures for *Acropora* spp.

We further detected variable effects of elevated temperature and oxybenzone on the microbiome of *S. pistillata*. The heat wave without oxybenzone treatment resulted in elevated abundance of the bacterial family Deinococcaceae. In contrast, oxybenzone exposure resulted in elevated abundance of Verrucomicrobiaceae, but not at elevated seawater temperature. Combined exposure to oxybenzone and elevated temperature resulted in decreased abundance of Hyphomonadaceae, Nocardioidaceae and Sinobacteraceae, as well as decreased abundance of 10 genera. The observed limited effects of temperature alone is in agreement with the study of Gardner et al. (2019), who found a conservative within-species microbial community in the Seychelles during a thermal bleaching event. Our study suggests that multistressor conditions, in our case elevated temperature and oxybenzone exposure, lead to a stronger shift within the coral microbiome. The significance of microbial changes in terms of coral health is not clear yet, although it is possible that many physiological and immunological processes within the coral holobiont are affected. The importance of the microbiome to coral health has been described by several authors, including its role in adapting to environmentally stressful situations (Gardner et al., 2019; Glasl et al., 2016; Peixoto et al., 2017; Ziegler et al., 2017; Grottoli et al., 2018). Indeed, environmental stress resulting in coral bleaching, tissue necrosis and mortality is often accompanied by a shift in the microbiome (Glasl et al., 2016; Zaneveld et al., 2017). Bulleri et al. (2018) found that coral recruitment is negatively correlated with the abundance of Verrucomicrobiaceae, and Closek et al. (2014) found Verrucomicrobiaceae exclusively in corals suffering from yellow band-disease. The fact that these bacteria are associated with negative conditions does however not make clear whether these bacteria cause coral disease or opportunistically increase in abundance due to impaired holobiont health. Whether the increase of Verrucomicrobiaceae in the oxybenzone treated corals benefited from an impaired holobiont, or had a steering role in the observed decreased health can thus not yet be elucidated. Generally, the various roles of coral-associated bacteria under healthy, stable conditions and in diseased states remains a critical knowledge gap. Functional gene analysis using metagenomics (Thurber et al., 2009) and transcriptomics (Miller et al., 2011), in combination with multifactorial laboratory experiments, may shed more light on how the coral holobiont responds to chronic stressors such as oxybenzone.

### Implications for risk assessment and management

For effective reef management, understanding the interplay between global and local stressors is vital (Ban et al., 2014). The identification of synergistic or added effects between stressors allows the prioritization of measures, such as improving local water quality to enhance resistance to thermal bleaching (Wooldridge, 2009; Carilli et al., 2010; Ban et al., 2014), thereby possibly creating a safer operating space for these ecosystems (Scheffer et al., 2015). Negri et al. (2011) showed that reducing the coastal herbicide (diuron) concentration by 1 µg L^−1^ in situations above 30 °C would increase the thermal tolerance of corals in terms of photosynthetic efficiency by the equivalent of 1-1.8 °C. Such results demonstrate the relevance of local water quality measures in order to give coral reefs more time to acclimate or adapt to climate change.

Taken together, our results support the current view that semi-chronic exposure to field-relevant levels of oxybenzone can cause subtle adverse effects on coral health, and a pronounced impact on the coral microbiome. Therefore, oxybenzone adds insult to injury by further weakening corals in the face of climate change. Future experiments should involve long-term oxybenzone exposure to assess cumulative effects, and should include more species, genotypes and life stages to better estimate the impact on coral reef health. In addition, a better understanding of the mechanisms through which oxybenzone affects corals, their zooxanthellae and their microbiome is important.

## Acknowledgements

The authors wish to thank Anna Haider Rubio, Sam Plaatsman, Nina Villing, Jochem van Herwaarden, Max Janse and Jorg Sander for their support. The project was financially supported through internal allocated funds by Wageningen Marine Research under project reference KB-24-002-039. There was no conflict of interest.

