## Supplementary figures and tables for "Adding insult to injury: Effects of chronic oxybenzone exposure and elevated temperature on two reef-building corals"

Additional figures (4) and tables (2) to manuscript:

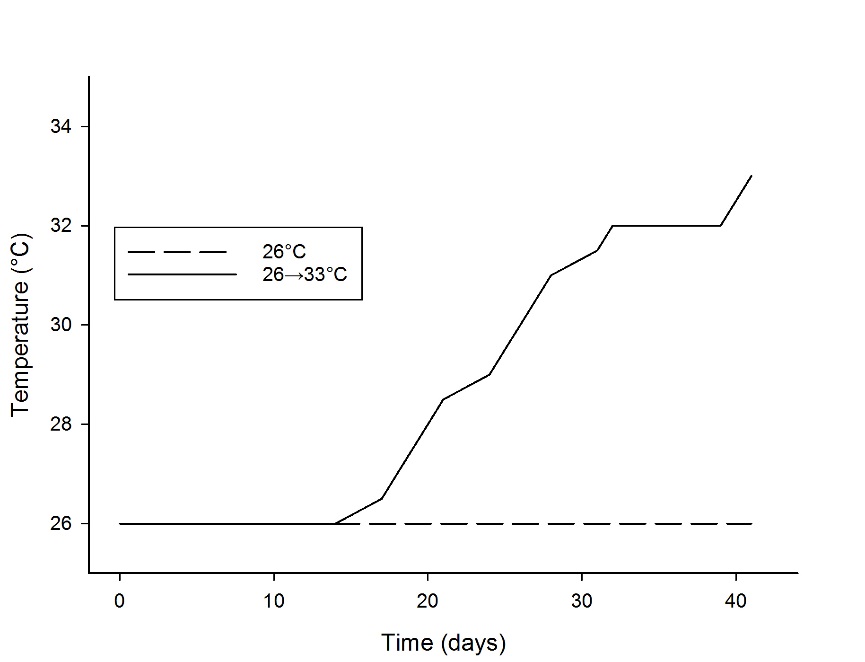

Figure S1. Overview of temperatures applied throughout the experiment (N=10 tanks per temperature group).

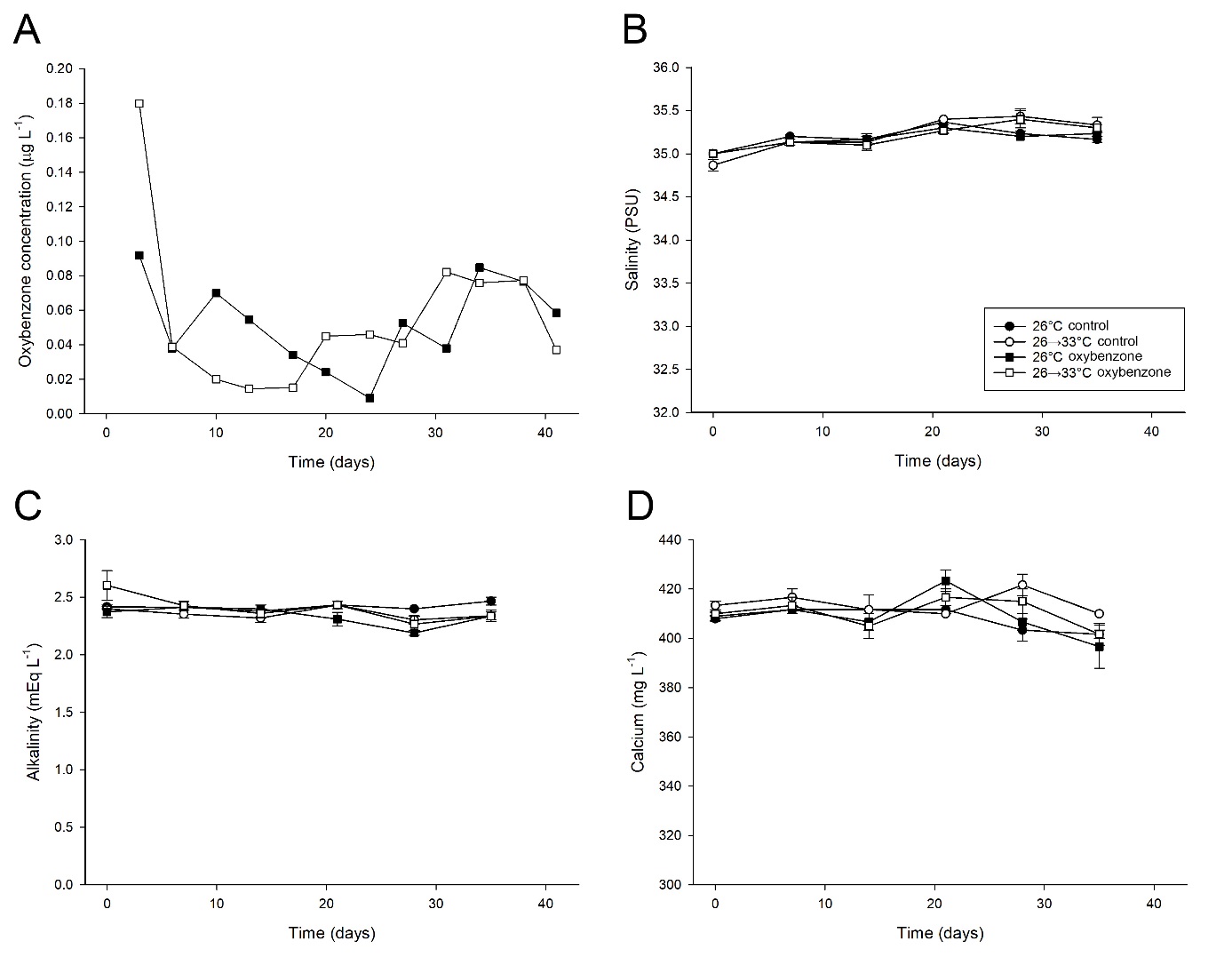

Figure S2. Overview of several water chemistry parameters during the experiment. A: recovered oxybenzone concentration of both temperature groups, B: salinity of all treatment groups, C: alkalinity of all treatment groups, D: calcium level of all treatment groups. A: Values are individual measurements, based on 5 pooled oxybenzone tanks for each temperature group, B-D: Values are means ± s.e.m. (N=3 tanks for each treatment group).

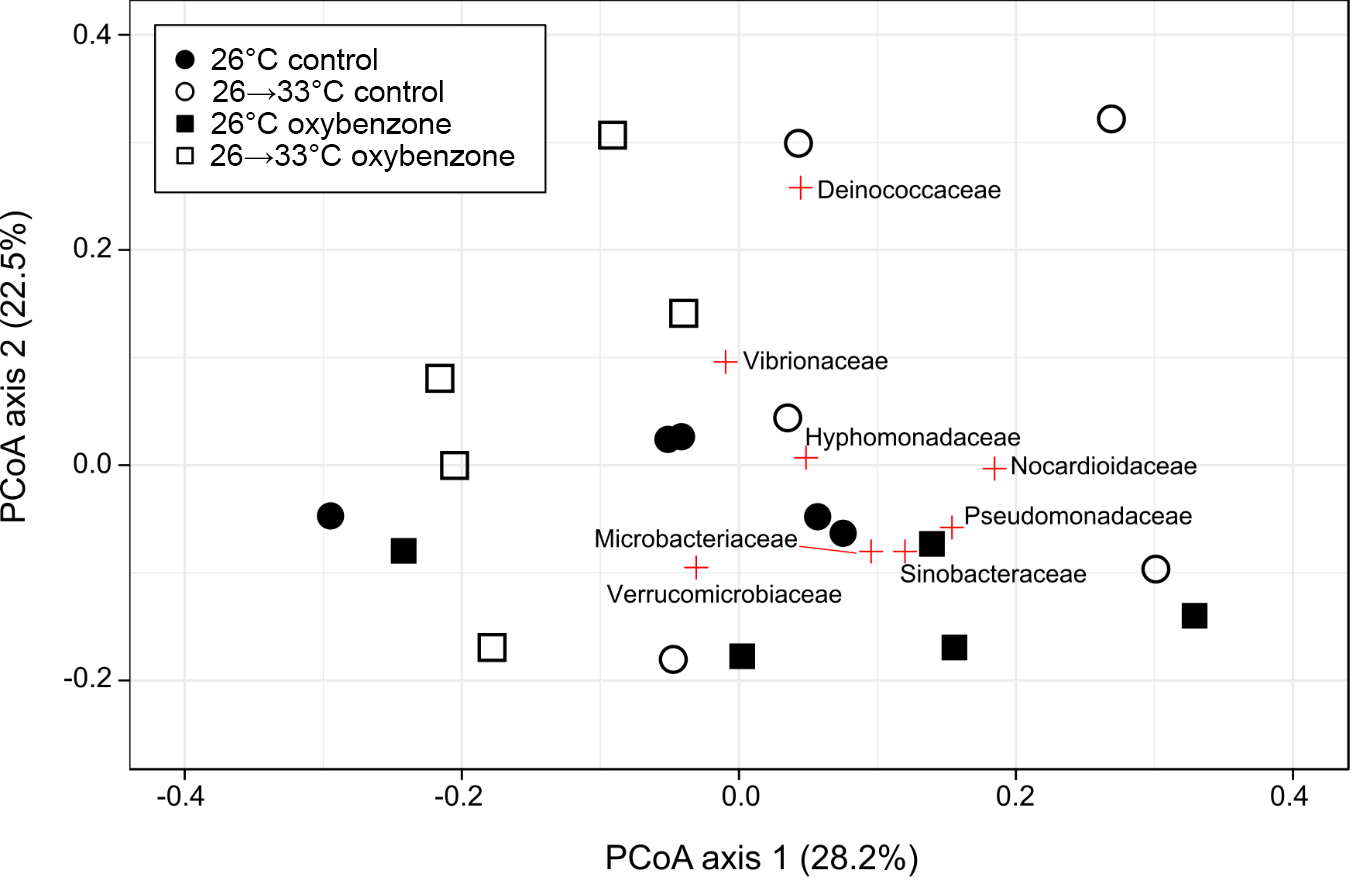

Figure S3. Principal Coordinates Analysis (PCoA) using a Bray-Curtis distance matrix of the *S. pistillata* microbiome composition. The PCoA was run on relative abundance of 16S bacterial DNA at family level. Families which explain most of the variability have been labelled in the plot.

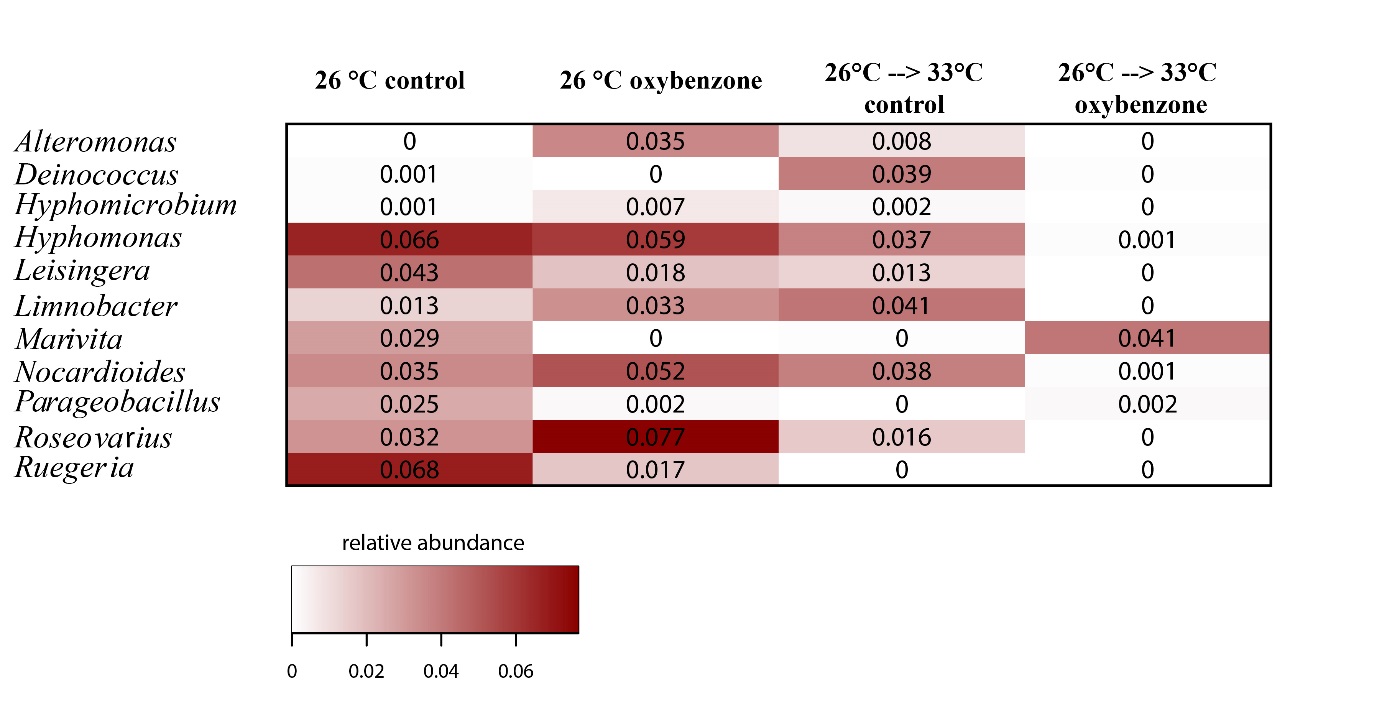

Figure S4. Relative abundance of genera that responded significantly to temperature and/or oxybenzone. Light colours indicate low relative abundance, dark red indicates high abundance.

Table S1. Overview of ANOVA’s performed on physiological data of *Stylophora pistillata* and *Acropora tenuis*. For effective and maximum PSII yield, a three-way mixed factorial ANOVA was used. For specific growth rate and zooxanthellae density, a two-way factorial ANOVA was used. See methods for details. *Sp*: *Stylophora pistillata*, *At*: *Acropora tenuis*. Empty cells indicate unavailable data. Asterisks indicate a significant effect, *: *p<*0.05, **: *p<*0.01, ***: *p<*0.001.

| Factor | Variable | F | | df | | *p* | |
| --- | --- | --- | --- | --- | --- | --- | --- |
|  | Effective PSII yield | *Sp* | *At* | *Sp* | *At* | *Sp* | *At* |
| Temperature |  | 37.62 |  | 1 |  | **0.000***** |  |
| Oxybenzone |  | 26.23 | 7.14 | 1 | 1 | **0.000***** | **0.037*** |
| Time |  | 26.04 | 2.38 | 25 | 25 | **0.000***** | **0.001**** |
| Temperature * Oxybenzone |  | 1.72 |  | 1 |  | 0.212 |  |
| Temperature * Time |  | 11.95 |  | 25 |  | **0.000***** |  |
| Oxybenzone * Time |  | 1.24 | 1.53 | 25 | 25 | 0.201 | 0.064 |
| Temperature * Oxybenzone * Time |  | 0.99 |  | 25 |  | 0.476 |  |
|  | Maximum PSII yield |  |  |  |  |  |  |
| Temperature |  | 33.55 |  | 1 |  | **0.000***** |  |
| Oxybenzone |  | 10.14 | 0.64 | 1 | 1 | **0.006**** | 0.448 |
| Time |  | 47.57 | 1.95 | 3.07 | 1.69 | **0.000***** | 0.184 |
| Temperature * Oxybenzone |  | 0.39 |  | 1 |  | 0.542 |  |
| Temperature * Time |  | 25.15 |  | 3.07 |  | **0.000***** |  |
| Oxybenzone * Time |  | 1.58 | 1.21 | 3.07 | 1.69 | 0.206 | 0.321 |
| Temperature * Oxybenzone * Time |  | 0.26 |  | 3.07 |  | 0.858 |  |
|  | Specific growth rate |  |  |  |  |  |  |
| Temperature |  | 36.15 |  | 1 |  | **0.000***** |  |
| Oxybenzone |  | 3.38 |  | 1 |  | 0.086 |  |
| Temperature * Oxybenzone |  | 0.02 |  | 1 |  | 0.895 |  |
|  | Zooxanthellae density |  |  |  |  |  |  |
| Temperature |  | 57.24 |  | 1 |  | **0.000***** |  |
| Oxybenzone |  | 2.60 |  | 1 |  | 0.128 |  |
| Temperature * Oxybenzone |  | 0.07 |  | 1 |  | 0.791 |  |

Table S2. PERMANOVA on microbiome beta diversity between oxybenzone and temperature treatments. Analyses are based on Bray-Curtis dissimilarity with fixed effects and 9999 permutations. Asterisks indicate a significant effect, *: *p<*0.05.

| Factor | Variable | Pseudo-F | df | *p* |
| --- | --- | --- | --- | --- |
|  | Order |  |  |  |
| Temperature |  | 2.40 | 1 | **0.01*** |
| Oxybenzone |  | 1.15 | 1 | 0.31 |
| Temperature * Oxybenzone |  | 2.12 | 1 | **0.03*** |
|  | Family |  |  |  |
| Temperature |  | 2.45 | 1 | **0.02*** |
| Oxybenzone |  | 0.99 | 1 | 0.43 |
| Temperature * Oxybenzone |  | 2.29 | 1 | **0.02*** |
|  | Genus |  |  |  |
| Temperature |  | 1.80 | 1 | 0.09 |
| Oxybenzone |  | 0.64 | 1 | 0.76 |
| Temperature * Oxybenzone |  | 2.43 | 1 | **0.03*** |
