## Supplementary information 1 for "Adding insult to injury: Effects of chronic oxybenzone exposure and elevated temperature on two reef-building corals"

Supplementary information 1. Oxybenzone analysis

*Sample extraction and analysis*

Samples were separated by solid phase extraction (SPE) using SEP-PAC VAC 6 cc (1 g) t-C18 cartridges. 100 ml of samples was weighed in a glass bottle. Samples were spiked with 200 µl D5-oxybenzone Internal standard solution (450 ng/ml in methanol). Cartridges were activated using 10 ml of acetonitrile followed by 20 ml of deionised water after which the sample was introduced. Samples were eluted using 20 ml acetonitrile and consequently concentrated to 1 ml using a Turbovap at 65 °C. Extracts were centrifuged and the top layer was transferred to a sample vial for analyses.

*Analysis by LC-MS*

Samples were analysed by LC-MS/MS. Separation was performed using an Agilent 1200 HPLC system containing an Agilent Zorbax Eclipse XDB-C18 2.1 x 150 mm 3.5 µm column coupled with a Micromass Quattro Micro MS instrument. Ammonium formate (0.02 mM) and formic acid (0.05 mM) in deionised water was used as mobile phase A and ammonium formate (0.02 mM) and formic acid (0.05 mM) in acetonitril as mobile phase B. Flow was held at 0.3 ml/min. Gradient started with 70% B, hold 1 min, 95% B at 1.1 min, hold 7.9 min, 70% B at 8.1 min, hold for 7 min. Quantification was performed in ESI negative mode using the following ions 229->151 and 229>105 as quantifier and qualifier for oxybenzone and 233.75->151 and 233.75>109.92 as quantifier and qualifier for D5-oxybenzone.
